# Selective estrogen receptor modulators limit alphavirus infection by targeting the viral capping enzyme nsP1

**DOI:** 10.1101/2021.08.19.457046

**Authors:** Rajat Mudgal, Chandrima Bharadwaj, Aakriti Dubey, Shweta Choudhary, Perumal Nagarajan, Megha Aggarwal, Yashika Ratra, Soumen Basak, Shailly Tomar

## Abstract

Alphaviruses cause animal or human diseases that are characterized by febrile illness, debilitating arthralgia, or encephalitis. Selective estrogen receptor modulators (SERMs), a class of FDA-approved drugs, have been shown to possess antiviral activities against multiple viruses, including Hepatitis C virus, Ebola virus, dengue virus, and vesicular stomatitis virus. Here, we evaluated three SERM compounds, namely 4-hydroxytamoxifen, tamoxifen, and clomifene, for plausible antiviral properties against two medically important alphaviruses, chikungunya virus (CHIKV) and Sindbis virus (SINV). In cell culture settings, these SERMs displayed potent activity against CHIKV and SINV at non-toxic concentrations with EC_50_ values ranging between 400 nM and 3.9 μM. Further studies indicated that these compounds inhibit a post-entry step of the alphavirus life cycle, while enzymatic assays involving purified recombinant proteins confirmed that these SERMs target the enzymatic activity of non-structural protein 1 (nsP1), the capping enzyme of alphaviruses. Finally, tamoxifen treatment restrained CHIKV growth in the infected mice and diminished musculoskeletal pathologies. Combining biochemical, cell culture-based studies, and *in vivo* analyses, we strongly argue that SERM compounds, or their derivatives, may provide for attractive therapeutic options against alphaviruses.

## Introduction

Alphaviruses are enveloped arboviruses and possess a positive-sense (+ve sense), single-stranded RNA genome. Symptomatic manifestations of alphavirus infection in humans vary from fever or rash to incapacitating polyarthralgia and intense asthenia persisting for months (1). These viruses can be broadly classified into New World alphaviruses, such as Venezuelan equine encephalomyelitis virus (VEEV), and Old World alphaviruses, such as chikungunya virus (CHIKV) and Sindbis virus (SINV). While Old World viruses mainly cause polyarthritis, New World viruses are predominantly associated with encephalitis in the infected host (2). In particular, numerous re-emergence cases of CHIKV infections were reported in the past 50 years; the most severe outbreak was observed in the year 2005 in La Réunion and parts of the Indian subcontinent resulting in significant morbidities, economic losses, and stress on the health care infrastructure (3, 4). On the other hand, SINV has been attributed to recurrent outbreaks in Scandinavia, Africa, Europe, and the Middle East, since 1952 (5). Despite significant human health relevance, there are currently no licensed antiviral drugs or vaccines to combat the alphavirus menace. Treatment of infection is palliative and consists of the administration of analgesics or antipyretics to impart symptomatic relief.

Post-infection, the alphaviral genome is translated to produce a non-structural and a structural polyprotein. Proteolytic cleavage of these polyproteins generates functional viral polypeptides (2, 6). Proteolytic cleavage of the non-structural polyprotein generates four non-structural proteins (nsPs) viz. nsP1, nsP2, nsP3, and nsP4, which play essential roles in viral RNA (vRNA) synthesis within host cells (7). In particular, nsP1, which is a membrane-associated enzyme, directs the capping of genomic and subgenomic vRNAs with its methyltransferase (MTase) and guanylyltransferase (GTase) activities (8). Capping of the vRNA is crucial for translation initiation, protecting vRNAs from host exonucleases, and masking viral nucleic acids from host-encoded pathogen recognition receptors, such as MDA5 and RIG-I (9). Due to a distinctive and non-canonical capping mechanism, nsP1 is thought to be a promising antiviral target for alphaviruses (10–12).

Identification and evaluation of novel antiviral therapeutics is a challenging and time-consuming process. However, repurposing drugs already in clinical use may expedite antiviral development against alphaviruses (13). 4-hydroxytamoxifen (4-OHT), tamoxifen, and clomifene belong to the triphenyl ethylene class of molecules and are known estrogen receptor antagonists (14, 15). Tamoxifen is typically used as a maintenance agent in estrogen-receptor-positive tumor patients (16). 4-OHT is a metabolite of tamoxifen (17). Clomifene is used for treating female infertility. Both tamoxifen and clomifene have also found their off-label use in treating male infertility and idiopathic gynecomastia (18, 19). These compounds, along with their analogs, are generally referred to as selective estrogen receptor modulators (SERMs) (15). Previous reports revealed that SERMs are active against multiple viruses, including the hepatitis C virus (HCV), herpes simplex type 1 virus (HSV-1), Ebola virus, dengue virus, vesicular stomatitis virus (VSV), etc (20–25). However, the plausible role of SERMs in limiting alphavirus infections has not been examined.

Here, we report potent antiviral activity of three SERMs, namely, 4-OHT, tamoxifen, and clomifene, against CHIKV and SINV in both cell culture-based settings and the preclinical mouse model. Our biochemical studies further established that SERMs inhibit the enzymatic activity of nsP1 from multiple alphaviruses. Taken together, SERM compounds may offer an effective means for antiviral therapy against the alphavirus genus.

## Materials and Methods

### Cells, animals, viruses, and compounds

Vero cells (NCCS, Pune, India) were maintained in DMEM (HiMedia, India) supplemented with 10% heat-inactivated FBS (Gibco), 100 units of penicillin, and 100 mg/ml of streptomycin at 37°C and in a 5% CO_2_ incubator. Stocks of CHIKV (accession number, KY057363.1; strain, 119067) and SINV were maintained using standard viral adsorption techniques (26, 27). (Z)-4-OHT, tamoxifen, and clomifene citrate were purchased from Cayman chemicals and dissolved in 100% dimethyl sulfoxide (DMSO). WT C57BL/6 mice were housed at the National Institute of Immunology (NII) and used adhering to the institutional guidelines (Approval number – IAEC 545/20).

#### *In vitro* antiviral assays

Cytotoxicity of various compounds was evaluated in MTT assay and LDH assay (see the supporting information). For inhibitor treatment of the infected cells, subconfluent Vero cells were infected for 1.5 h at MOI:1 in a 24-well plate at a density of 1×10^5^ cells/well. The media was then removed, and compound dilutions, lower than their respective toxic doses, prepared in low-serum media were added and cells were incubated at 37°c. At 24 h post-infection (hpi), culture supernatant was harvested and subjected to plaque-forming assay (see a detailed description of the plaque assay in the supporting information). The reduction in viral titer relative to DMSO-treated cells was measured and graphs were analysed using Graph Pad Prism for determining the half-maximal effective concentration (EC_50_) values. The antiviral activity of various compounds was also examined in the indirect immunofluorescence assay, where intracellular viral particles were stained using anti-CHIKV antibodies (see the supporting information for details). Cells were harvested at various time points and the abundance of intracellular and extracellular vRNA was quantified in the qRT-PCR assay. A detailed description of the luciferase-based antiviral assays, qRT-PCR protocol, and primers are available in the supporting information.

#### nsP1 expression, purification, and enzymatic assays

CHIKV nsP1 and VEEV nsP1 were expressed and purified from *E. coli* as described in a previous study (12). Reaction conditions and data analysis for capillary electrophoresis (CE)-based MTase assay and enzyme-linked immunosorbent assay (ELISA)-based guanylation assay were described earlier (12, 28).

#### CHIKV infection in mice and tamoxifen treatment

Six weeks old male mice were subcutaneously (SC) injected on the ventral side of the right footpad with 10^6^ PFU of CHIKV (strain no: 119067) (29). CHIKV-infected mice were injected with tamoxifen dissolved in sesame oil at day+1, day+3, and day+5 post-infection. Control mice were injected with an equal volume of oil. Infected mice were further examined for body weight changes at the indicated days post-infection. The swelling of the infected footpad was quantified by estimating the increase in the footpad upon infection using Insize digital Vernier caliper (30). In certain sets, mice were sacrificed at day six, and tissues from decalcified footpads were subjected to H&E staining for scoring pathologies associated with musculoskeletal inflammation. At the indicated days post-infection, blood was collected through the retro-orbital route, and the footpad was dissected from the infected mice. Total RNA was isolated from blood or tissue samples and the abundance of vRNA was determined by qRT-PCR using primers specific for the +ve sense genome (31). Separately, we prepared a serial dilution of CHIKV stock, whose titer was determined using the standard plaque assay. RNA was isolated and qRT-PCR was performed using these serial diluents. By plotting C_T_ values against the known CHIKV titer, a standard curve was generated. We used this standard curve for estimating the number of copies of RNA genomes present in samples from infected mice. Alternatively, the viral titer in these tissues was determined by the TCID_50_ method (see supporting information) (32).

## Results

### SERMs inhibit alphaviral propagation

As SERMs were active against multiple viruses, we sought to examine if this family of drugs also display antiviral properties against alphaviruses. Prior to the evaluation of antiviral activity, cell viability of BHK-21 and Vero cells in the presence of compounds was determined by MTT [3-(4,5-Dimethylthiazol-2-yl)-2,5-Diphenyltetrazolium Bromide] assay (Figure S1A-C and S2A-C). An additional cytotoxicity assay, Lactate Dehydrogenase assay (LDH) assay, which assesses the membrane integrity of cells, was also performed in Vero cells (Figure S2D-F). The calculated half-maximal cytotoxic concentration (CC_50_) values are presented in Table 1. A luciferase reporter system, SINV-FLuc, that has a *firefly luciferase* gene fused at the 3’end of the SINV E1, was used for primary evaluation of antiviral activity of compounds (Figure 1A). The compounds were tested in BHK-21 cells at 2.5 μM concentration at which there was no observable cytotoxicity. All the three compounds showed a significant decrease in luciferase activity that was used as a surrogate of viral replication in the infected cells (Figure 1A).

**Figure 1:**
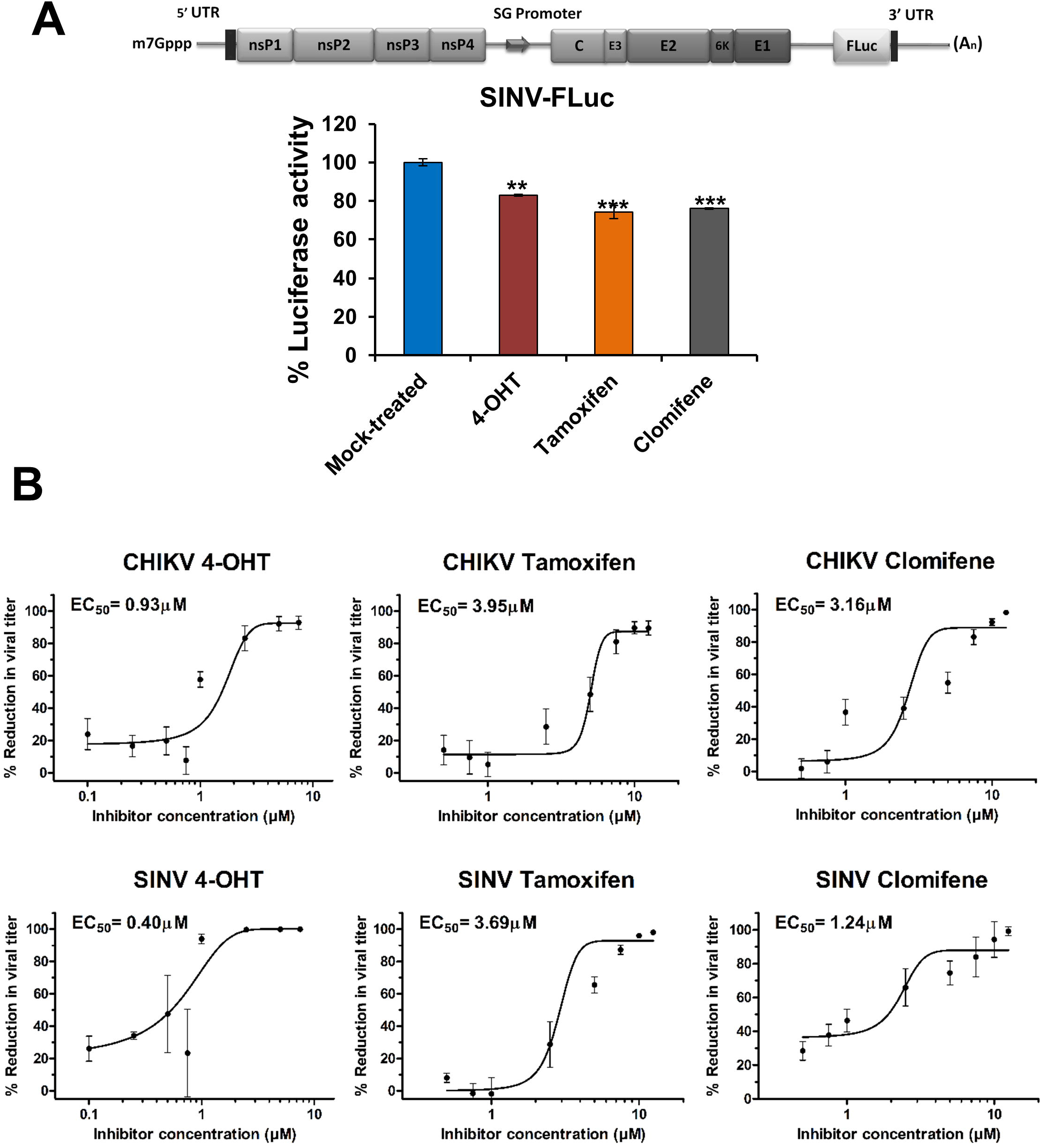
Primary evaluation of anti-alphaviral activity of SERMs. (A) Schematic diagram of SINV-FLuc (top panel) wherein FLuc is placed downstream of an internal ribosomal entry site (IRES). BHK-21 were infected with SINV-FLuc at an MOI of 0.1 and treated with SERM compounds or 0.1% DMSO. Luciferase activity of the cell lysate was measured 12 hpi using firefly luciferase assay kit. Values are mean and error bars are standard deviation (n=3). FLuc: Firefly luciferase; hpi: hours post-infection. Statistical significance of the difference in luciferase activity levels between treated and vehicle control-treated cells was assessed by one-way ANOVA test and Dunnett’s posttest. ***, p<0.001; **, p<0.01. Values are mean and error bars are standard deviation (n=3). (B) Plaque assay revealing changes in CHIKV (top three panels) or SINV (bottom three panels) titer on the treatment of virus-infected cells with an increasing concentration of 4-OHT, tamoxifen, and clomifene. The viral titer in the culture supernatant was determined at 24 hpi. The viral titer in corresponding vehicle-treated cells was considered 100%. Data were normalized using Graph pad’s non-linear regression curve fit and the calculated 50% effective concentration (EC_50_) values are summarized in Table 1. Values are mean from three independent experiments and error bars are standard deviation.

**Table 1:**
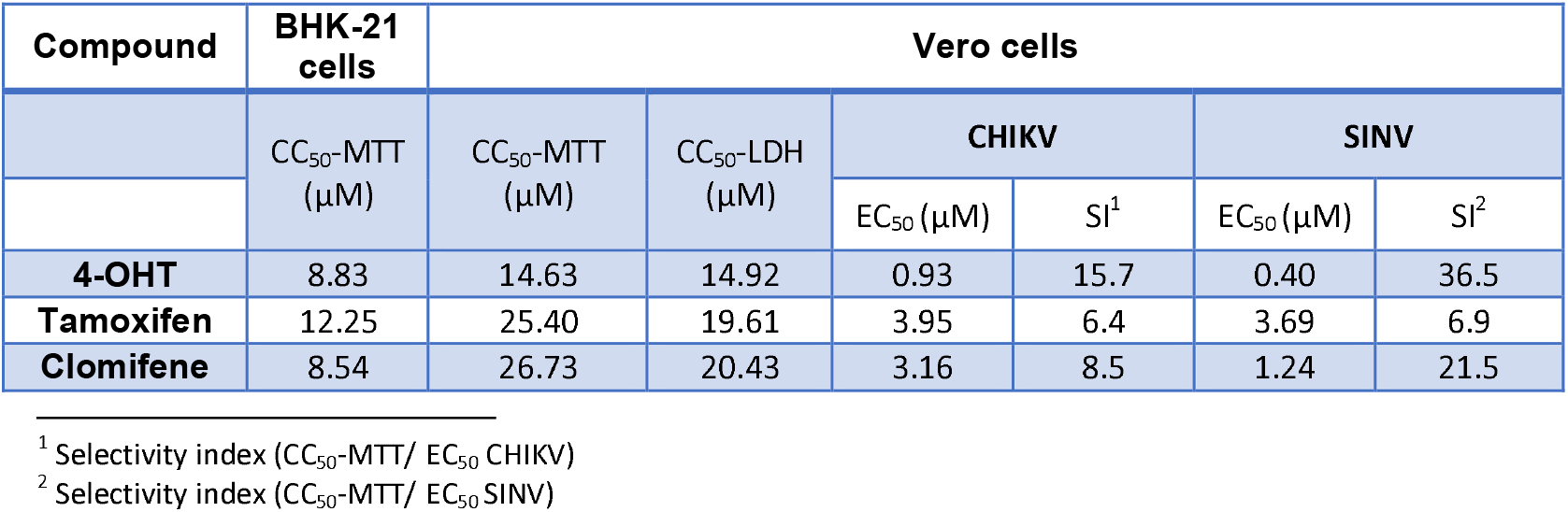
The CC50, EC_50_, and selectivity index values of 4-OHT, tamoxifen, and clomifene in BHK-21 and Vero cells.

The quantitative analysis of the antiviral activity of the compounds was done by the titer determination of the progeny virus released 24 hpi in the supernatant of the treated or untreated cells infected with CHIKV or SINV. All three compounds demonstrated a marked reduction in infectious virus production in a dose-dependent manner (Figure 1B). CHIKV-infected Vero cells treated with 4-OHT or tamoxifen showed maximally 1 log_10_ reduction relative to mock-treated cells while clomifene treatment reduced viral titer by 2 log_10_ at higher concentrations. Similarly, SINV titers were reduced maximally by 2 log_10_ when infected cells were treated with clomifene or tamoxifen. Even greater viral titer reduction (>3 log_10_) was observed upon 4-OHT treatment of SINV-infected Vero cells. Selectivity index values calculated from EC_50_ and CC_50_ suggested that within an order of magnitude, these drugs displayed largely comparable selectivity against CHIKV and SINV (Table 1). The anti-CHIKV and anti-SINV activities of 4-OHT, clomifene, and tamoxifen were further validated by qRT-PCR. There was a significant reduction in intracellular vRNA in the cells treated with the tested compounds 24 hpi. (Figure 2A). Corroborating our plaque assay, immunostaining cells with an anti-alphavirus antibody confirmed that treatment of CHIKV- or SINV-infected cells with 4-OHT, tamoxifen, or clomifene substantially suppress the abundance of viral proteins at 36 hpi (Figure 2B).

**Figure 2:**
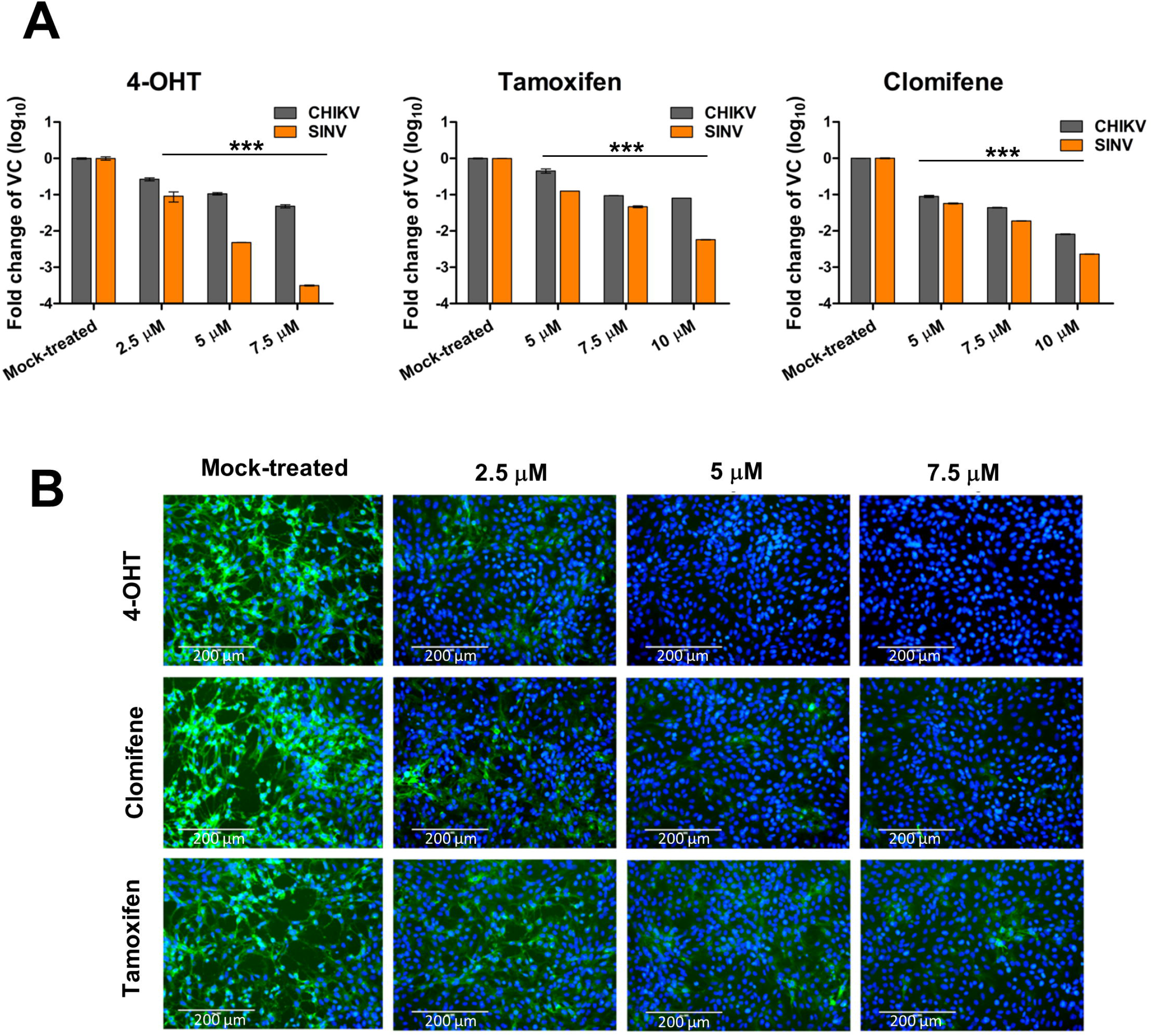
Antiviral activity of SERMs against CHIKV and SINV in cell culture. (A) Barplots showing fold-reduction in the abundance of vRNA in CHIKV or SINV infected Vero cells treated with SERMs compared to vehicle control (VC). At 24 hpi, RNA was isolated and analyzed by qRT-PCR for the expression of the E1 region of the viral genome. Statistical significance of the difference between intracellular vRNA in treated cells and vehicle control-treated cells was assessed by one-way ANOVA test and Dunnett’s posttest.***, p<0.001. Values are mean and error bars are standard deviation (n=3). (B) Immunofluorescence microscopy of CHIKV infection in Vero cells upon treatment with SERMs. Virus infection and treatment with SERMs were performed as mentioned in the supporting information. At 36 hpi, the infected cells were fixed and stained using an anti-alphavirus primary antibody and FITC-conjugated secondary antibody. DAPI was used for counter-staining cell nuclei. The image represents three independent biological replicates. Bars, 200 μM.

### Inhibition of alphaviral replication at a post-entry step

To gather insight into the mode of action of these compounds, time-of-addition studies were performed (Figure 3A-D). When cells were pretreated with the compounds before virus infection, minimal inhibitory effect on the viral yield was observed. A cotreatment assay was performed to examine if the compounds can affect viral binding and entry into the cell. Viral yield remained largely unaffected when compounds were present during infection suggesting that compounds do not interfere with virus particle stability or its attachment to the host cell. Moderate dose-dependent inhibition of viral replication observed in cotreatment assay can be attributed to the possible effects of the residual amount of inhibitors retained by the cells during cotreatment (Figure 3B). All three compounds were most effective against both CHIKV and SINV when they were added during an early post-entry phase of viral replication in the infected cells suggesting a similar antiviral mechanism of the inhibitors in these two alphaviruses (Figure 3C-D).

**Figure 3:**
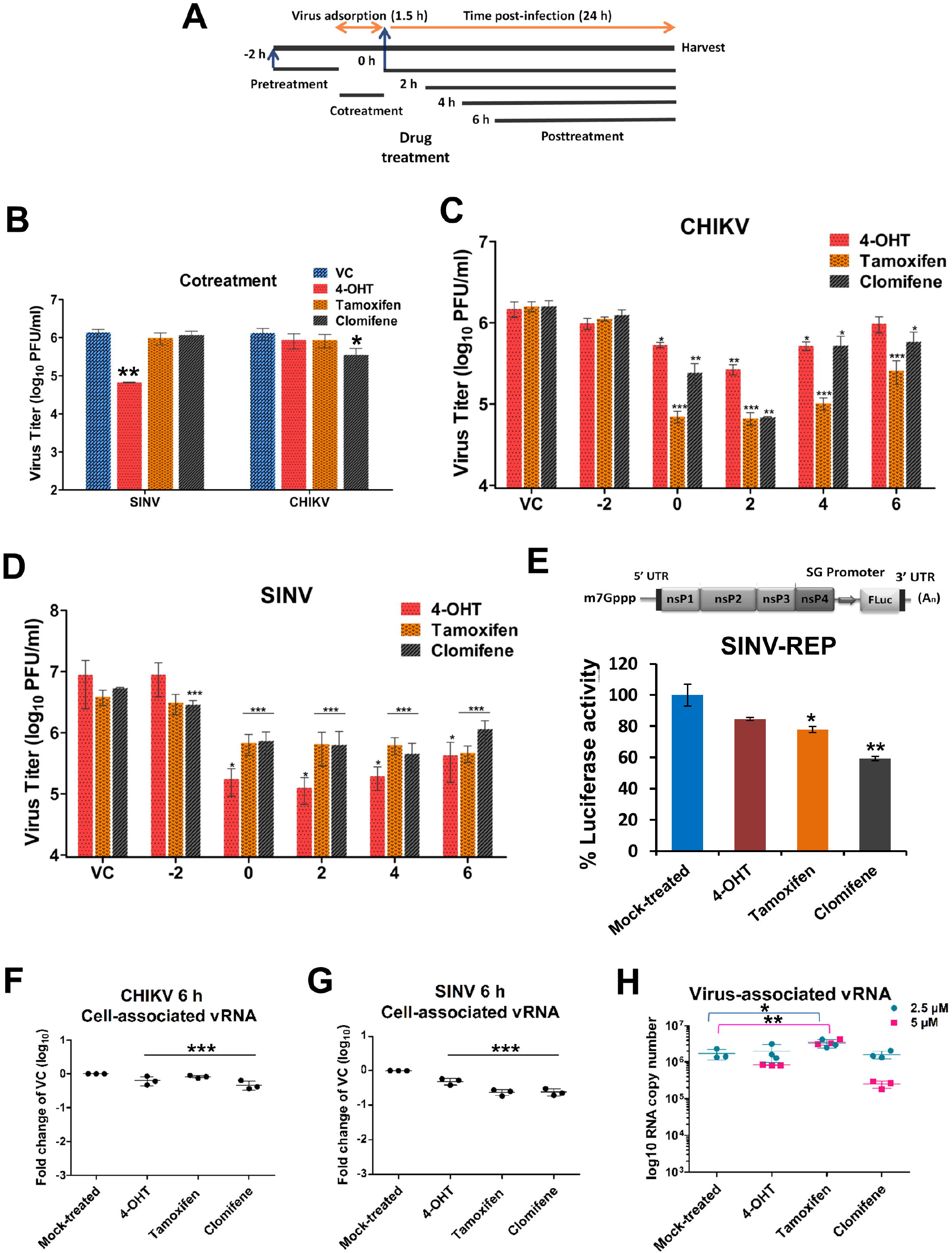
Characterizing SERM-mediated inhibition of viral growth. (A) Schematic representation of the time course of viral infection and treatment with SERM compounds. Vero cells were treated with compounds at different time points (in hours) before, during, or after virus infection and the viral titer (PFU/ml) was determined for CHIKV (B and C) and SINV (B and D). For Cotreatment studies, compounds were added simultaneously with the virus inoculum. For pre-treatment assay, Vero cells were treated with the inhibitors 2 h prior to viral adsorption. For post-treatment assay, cells were incubated at 37°C with viral infection medium which was replaced by maintenance medium at different time-points containing 5 μM 4-OHT or 7.5 μM tamoxifen/clomifene. Supernatants were collected 24 hpi for the quantification of viral titer by plaque assay. 0.1% DMSO was used as vehicle control. For all the experimental groups, Vero cells were infected with CHIKV or SINV at an MOI of 1. VC: vehicle control; hpi: hours post-infection; PFU: plaque-forming unit. (E) SINV minigenome replicon assay capturing the luciferase activity in lysate derived from BHK21 cells transfected with capped RNA encoding the SINV-REP reporter replicon. Cells were subjected to 6 h of treatment with VC or 2.5 μM of the indicated SERMs. A schematic diagram of SINV-REP has also been presented (top). (F and G) Changes in vRNA levels in CHIKV and SINV infected cells at 6 hpi. Vero cells were infected with CHIKV and SINV at MOI of 5 followed by treatment with 5 μM of the compounds. vRNA levels were assessed by qRT-PCR and the fold reduction in vRNA relative to VC-treated cells is plotted. Actin: endogenous control; E1: target gene. (H) Quantitative determination of extracellular SINV vRNA levels at 24 hpi. Vero cells were infected with SINV at MOI of 1 and were treated with indicated concentrations of the SERM compounds. Quantification of the vRNA in the culture supernatant was done using qRT-PCR as discussed in the material and methods section. Statistical significance of the difference in viral titer/luciferase activity/vRNA levels between treated and vehicle control-treated cells was assessed by one-way ANOVA test and Dunnett’s posttest. ***, p<0.001; **, p<0.01; *, p<0.05. Values are mean and error bars are standard deviation from triplicate experiments.

A SINV reporter replicon was designed to investigate the inhibitory effect of the compounds (Figure 3E). The tested compounds showed a moderate inhibitory effect against SINV replicon indicating a weak suppression of synthesis of viral genome and proteins by nsPs (Figure 3E). Additionally, to examine if the compounds exert an inhibitory effect on vRNA replication and to minimize the contribution of multiple rounds of viral infection, intracellular vRNA from virus-infected cells was quantified at 6 hpi. A statistically significant but modest decrease in cell-associated vRNA was observed 6 hpi for CHIKV and SINV (Figure 3F and Figure 3G). To further examine the mechanistic function of these SERMs against alphaviruses, SINV vRNA in the extracellular fractions was quantified (Figure 3H). Intriguingly, there was markedly less reduction in extracellular virus-associated vRNA as compared to the infectious virions released at 24 hpi (Figure 3H and Figure 1B).

Further, CHIKV was serially passaged in the presence of gradually increasing concentrations of compounds. Viral-induced cytopathic effects (CPE) were monitored and no change in viral susceptibility to the inhibitors was observed even after multiple passages. Only partial resistance to 4-OHT could be observed after 15 passages which suggested that a high resistance is indeed difficult to generate against these compounds. Plaque-purified variant harboring all four compensatory mutations also displayed an altered plaque phenotype compared to the wild-type virus but did not offer resistance against tamoxifen or clomifene (Figure 4A-B). This partially-resistant variant displayed ~4-fold resistance (EC_50_ 4-OHT-resistance variant [3.99 μm]/EC_50_ wild-type [0.93 μM]) relative to wild-type virus (Figure S3) The sequencing data of partially-resistant isolates revealed four non-synonymous mutations in the nsP-coding region (Figure 4B). Two mutations lie in the zinc-binding and macro domains of the nsP3-coding region, one is in disordered region of the nsP4-coding region, and one lies in the MTase/GTase domain of nsP1. While the mutations in nsP3 and nsP4 fall in the enzymatically-inactive regions, the point of mutation in nsP1 lies in the catalytic domain of the enzyme indicating that the likely molecular target of SERM compounds is alphaviral nsP1. Additionally, mutations identified within nsP3 and nsP4 proteins correlate with domains of the proteins that are known to, or have strong evidence for, the protein-protein interaction (33–35).

**Figure 4:**
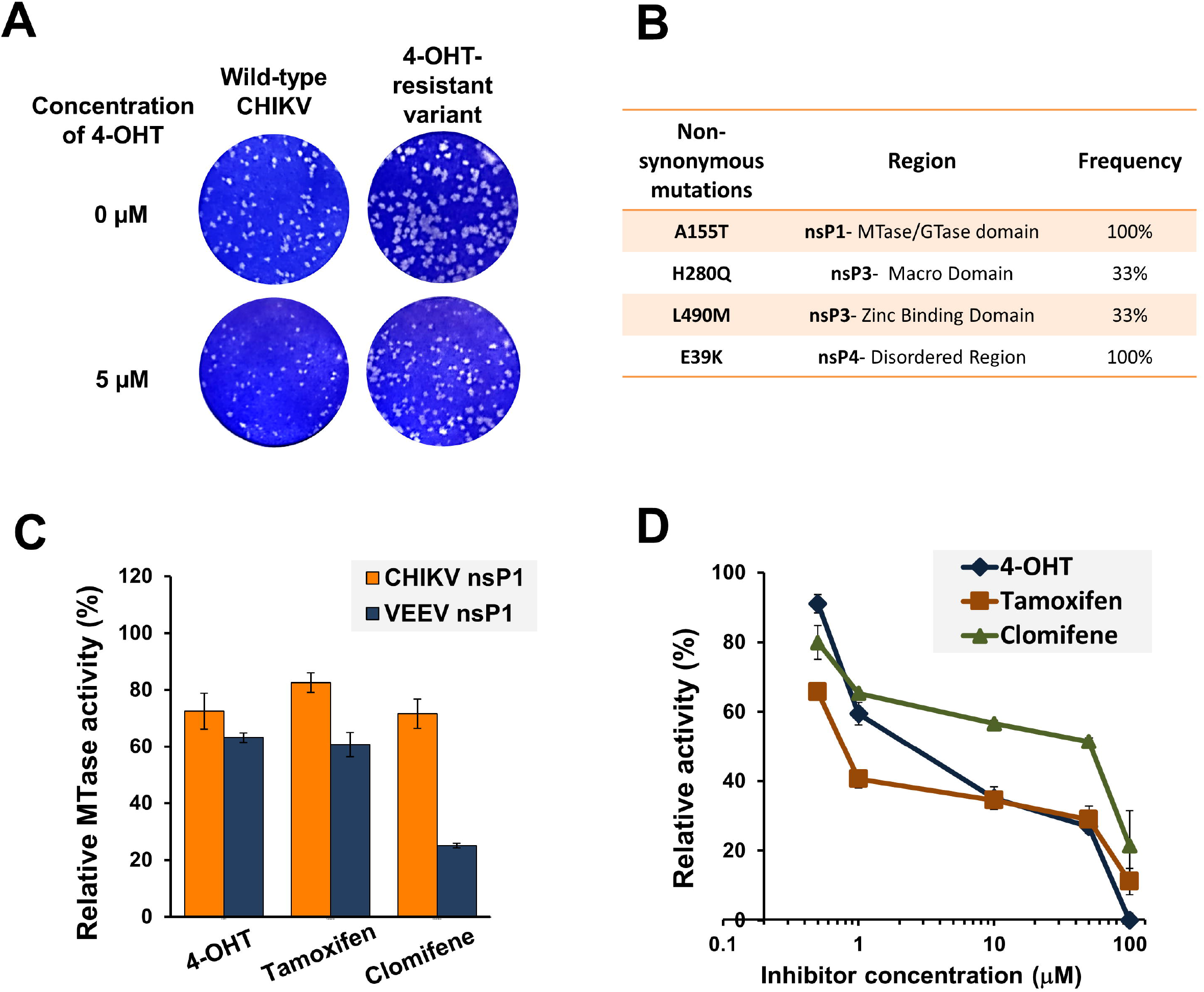
nsP1 capping enzyme is implicated in alphavirus inhibition by SERMs. Altered plaque phenotype of the partially-resistant variant of CHIKV against 4-OHT. (A) Standard plaque assay performed on the indicated virus. Vero cells were overlaid with 1% carboxymethyl cellulose-containing medium either without or with 5 μM 4-OHT. Plaques were visualised 48 hpi using crystal violet. Plaque-purified variant harboring all four mutations was used for phenotypic characterization. The resistant variant replicates more efficiently than wild-type CHIKV both in terms of number and size of plaques in the presence of 4-OHT. (B) Non-synonymous mutations in the nsP-coding region of partially-resistant CHIKV variant. Mutations were identified with the sequencing of three independent plaque-purified viruses. nsP: non-structural proteins. (C) nsP1 MTase activity inhibition by tested compounds. SERM compounds were tested at 50 μM concentration in the CHIKV nsP1 (10 μM) and VEEV nsP1 (20 μM) MTase reaction mixture. The reaction was carried out as described previously by Mudgal et al., 2020. The MTase activity is shown relative to the enzyme activity in the reaction mixture in the presence of vehicle control. (D) Dose-dependent inhibition of guanylation of nsP1 in the presence of SERMs. Relative guanylation by CHIKV nsP1 was determined using an ELISA assay that measures the amount of m^7^GMP-nsP1 adduct formed in the reaction. Reaction conditions were similar to those described earlier by Kaur et al., 2018. Data points represent mean value and error bars are standard deviation from two independent experiments.

### SERM compounds inhibit alphaviral nsP1 activity

Results of the studies described above prompted us to evaluate the effect of these compounds on the enzymatic activity of alphaviral nsP1. A non-radioactive CE-based MTase assay that measures the amount of SAH generated during CHIKV and VEEV nsP1 MTase reactions was utilized (12). Further, an ELISA-based assay that measures the m^7^GMP-nsP1 adduct formed in guanylation reaction was used to determine the inhibitory activity of compounds on CHIKV nsP1 (28). We observed that the compounds inhibited the MTase activity of both CHIKV and VEEV (Figure 4C). The ELISA-based assay also showed a concentration-dependent reduction in CHIKV nsP1 enzymatic activity in the presence of tested SERMs (Figure 4D).

### Tamoxifen alleviates CHIKV-induced pathologies *in vivo*

Next, we set out to examine if the anti-alphaviral property of SERMs was preserved *in vivo.* To this end, we adopted the previously published CHIKV footpad infection model involving adult wild-type mice that recapitulates certain arthritogenic phenotypes observed in human subjects (29). As described earlier, subcutaneous CHIKV infection into the footpad caused footpad swelling peaking at day two and again at day six (Figure S4) (29). Using tamoxifen as a representative drug, we examined the efficacy of SERMs in reducing these inflammatory musculoskeletal pathologies. We optimized a therapeutic regime where CHIKV-infected mice were successively injected with 20 mg/kg of tamoxifen on every alternate day beginning at day one post-infection. We found that compared to the vehicle control, tamoxifen treatment significantly improved the footpad swelling at day six post-infection, while somewhat moderately restraining footpad pathology at day two (Figure 5A-B).

**Figure 5:**
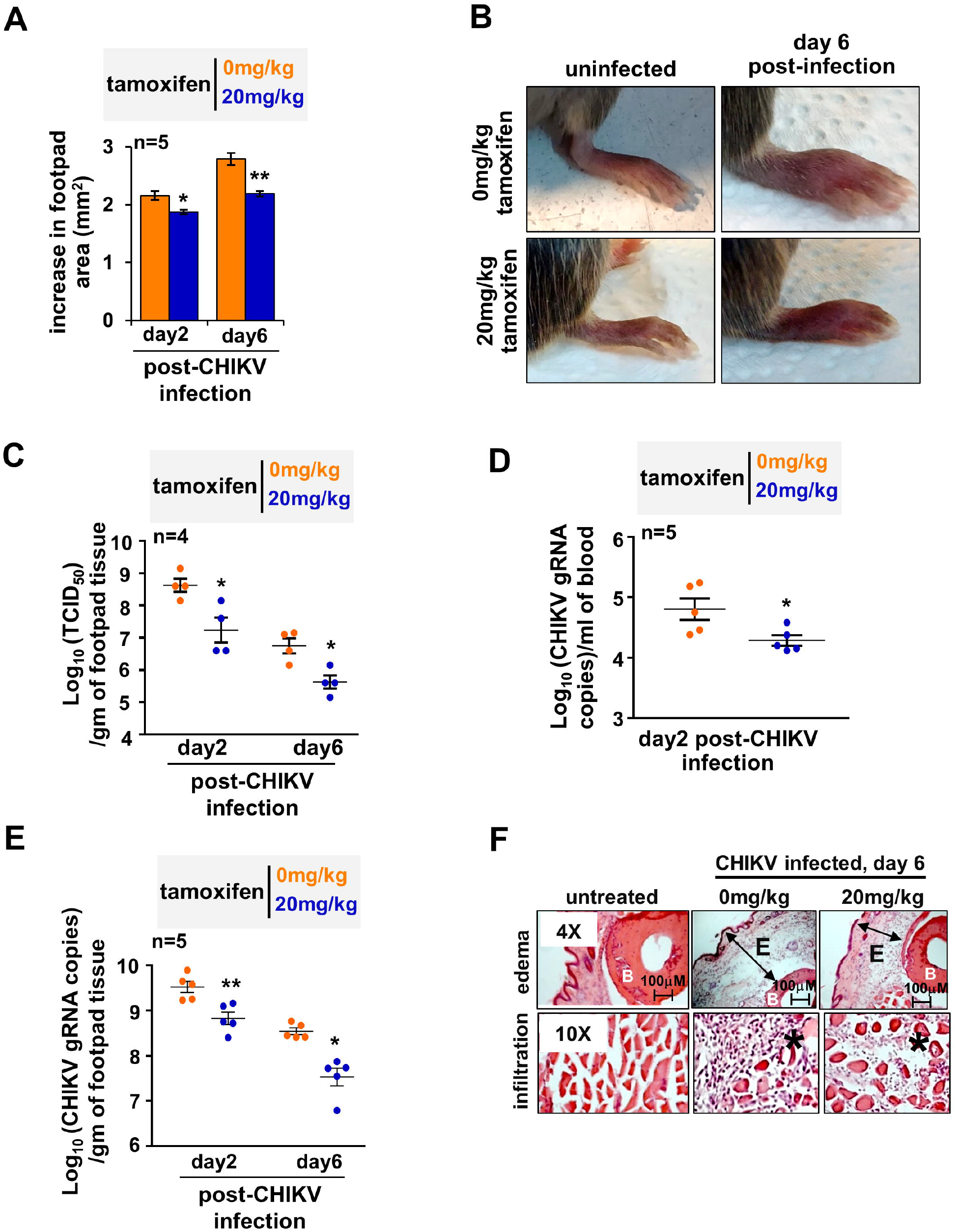
Testing therapeutic efficacy of tamoxifen in CHIKV-infected mice. CHIKV-infected mice were injected successively with 20 mg/kg of tamoxifen or vehicle control at day one, three, and five post-infection. Subsequently, changes in the footpad area (A) was measured at the indicated days. (B) Representative image of CHIKV infected joint footpad of mice subjected to tamoxifen therapeutic regime. (C) Dot plots showing the load of infectious virus particles in the footpad at the indicated days post-infection, as determined by TCID_50_ method. Graphs revealing the abundance of vRNA in the blood (D) and the infected footpad (E) at the indicated day post-infection. (F) Representative immunohistochemistry images revealing H&E-stained sections of the joint footpad. Mice were either uninfected (left panel) or infected with CHIKV six days prior to image acquisition (middle panel) or infected similarly with CHIKV and then subjected to a therapeutic regime (right panel). Footpad sections from two animals per set and three fields/sections were examined. E denotes edema; * denotes infiltrates; B signifies bone. Images are representative of three mice per group. Data were quantified from the indicated number of mice from two independent experimental replicates and presented as mean ± SEM. Two-tailed Student’s t-test was performed to assess statistical significance. **P<0.01; *P<0.05.

Because tamoxifen also directly impedes inflammatory pathways, we asked if the curative tamoxifen effect was indeed associated with reduced viremia *in vivo* (36). Consistent with the reduced musculoskeletal pathology, tamoxifen treatment led to a more than one log decrease in the titer of plaque-forming CHIKV in the infected footpad at both day two and day six post-infection (Figure 5C). Similarly, we captured an approximately five-fold decrease in the level of vRNA in the blood of tamoxifen-treated mice at day two (Figure 5D). In parallel, tamoxifen treatment ensured a close to one log reduction in the level of viral genome RNA in the footpad at both day two and day six (Figure 5E). Histopathological analyses of tissues derived from the joint footpad revealed that CHIKV infection triggered marked subcutaneous edema, profound infiltration of mononuclear immune cells, and local necrosis at day six (Figure 5F) (29). Tamoxifen treatment regime considerably ameliorated these signatures of CHIKV-induced inflammation. In sum, our analyses confirmed that SERMs are capable of restraining viremia and mitigating immunopathologies in alphavirus-infected mice.

## Discussion

Based on the antiviral effect exhibited by multiple SERMs against other virus families, we attempted to characterize the anti-alphaviral activity of three SERMs: 4-OHT, tamoxifen, and clomifene. As evident by plaque reduction assay, RT-PCR studies, and immunostaining, all these three compounds effectively abrogated the replication of SINV and CHIKV in cell culture. Results of the time-of addition studies and sequence analysis of partial-4-OHT resistant phenotype suggested that SERM compounds likely target nsP1 capping enzyme, a component of the vRNA replication machinery. In the presence of SERMs, a marked difference was observed between the number of infectious virions released and extracellular vRNA. Capping of vRNA in the infected cells is orchestrated by a functional alphaviral nsP1 and inhibition of nsP1 by SERM compounds would release a significant number of noncapped virus particles. The discrepancy in the magnitude of effect on extracellular vRNA and infectious units released is supported by previous studies which demonstrated that noncapped vRNAs are infectious but at significantly lower levels as compared to the capped RNAs (Figure 6) (37, 38). Though, SINV RNA has been shown to undergo translation and encapsidation without a cap structure, noncapped vRNAs are found to be highly unstable (38). Concordantly, the enzymatic activity of CHIKV nsP1 and VEEV nsP1 was inhibited by all the three compounds tested in the MTase activity assay. A secondary orthogonal ELISA-based assay also validated the anti-nsP1 activity of the tested SERM compounds.

**Figure 6:**
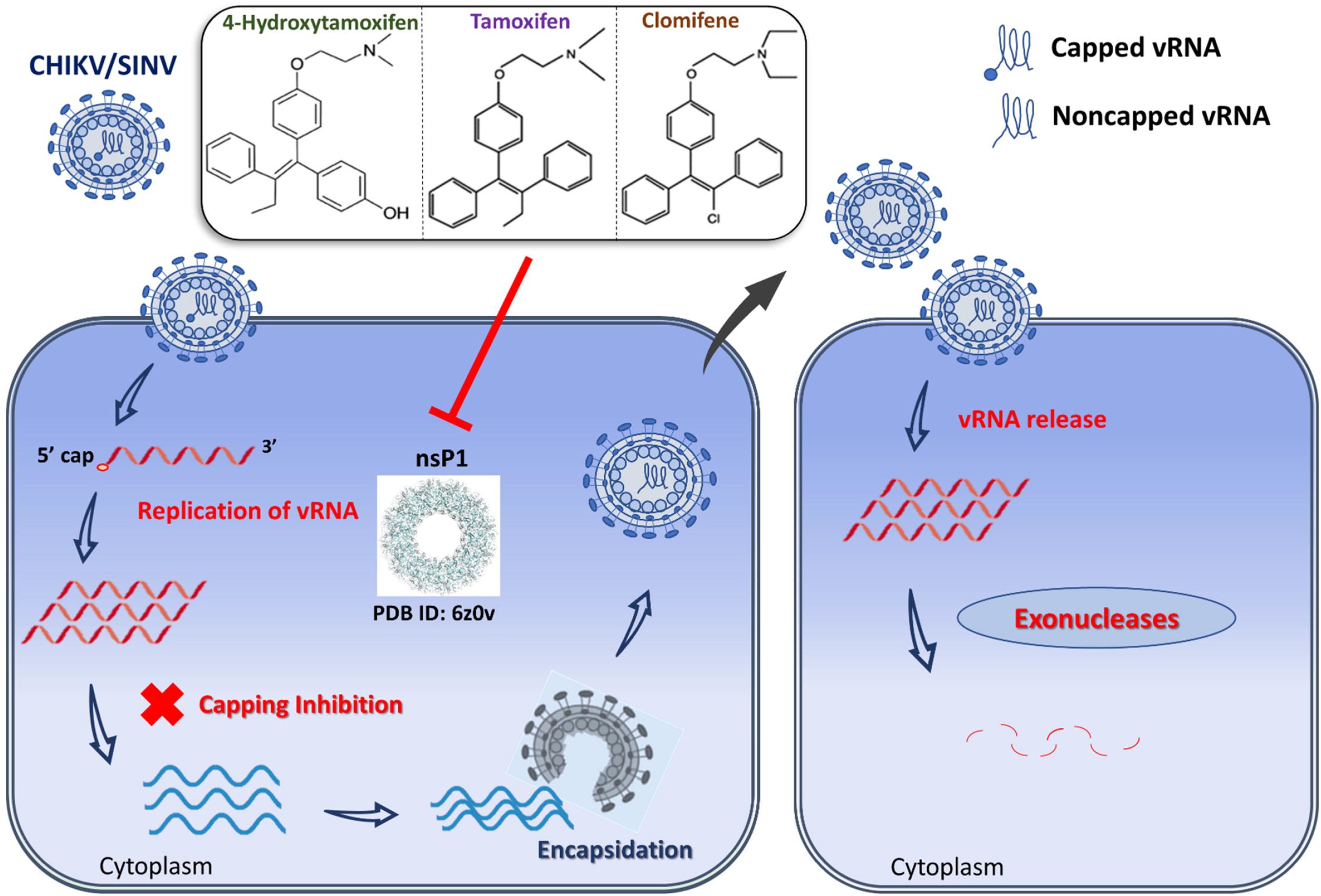
Possible mechanism of anti-alphaviral action of the SERM compounds mediated via nsP1 inhibition. vRNA is released in the cytoplasm upon entry of SINV/CHIKV in the infected cell. Subsequent translation of viral replication proteins generates viral RNA synthetic complex that in turn synthesizes numerous copies of vRNA. nsP1 (PDB ID: 6z0v) plays a critical role in 5ū capping of progeny genome. Putative inhibition of nsP1 by SERM compounds viz. 4-OHT, tamoxifen, and clomifene produces significant amount of noncapped nascent vRNAs that are encapsidated and subsequently released as mature virus particles. Upon entry into the host cell cytoplasm, these noncapped vRNAs undergo degradation by host RNA decay machinery following nucleocapsid disassembly and vRNA release.

Replicon assay using SINV-REP which does not involve viral entry or release indicated that these SERMs interfere with the viral replication machinery and subsequently affect viral protein synthesis. The data from the SINV-FLuc virus also suggested a decrease in vRNA replication and/or translation when the infected cells were treated with these compounds. The observed decrease in RNA translation can be a direct consequence of the reduction in capping activity by nsP1 or a result of perturbation of RNA replication machinery due to inhibitor binding to nsP1. We further demonstrated that these compounds marginally suppress vRNA replication in infected cells when vRNA was quantified 6 hpi. Apart from vRNA capping, nsP1 has also been found to be involved in the synthesis of negative strand of vRNA (39, 40). However, we cannot rule out the possibility that reduction of vRNA levels or subgenomic translation is due to the degradation of noncapped vRNAs (generated because of the compound-mediated inhibition of nsP1) by host RNA decay machinery or plausible modulation of nsP1 interaction with other nsPs due to inhibitor binding.

Besides engaging estrogen receptor signaling, SERMs also modulate numerous other cellular factors and functions; they inhibit immune-regulatory NF-κB signaling, repress the protein kinase C activity, control the calmodulin pathway, and restrict acidification of lysosomes and endosomes (36, 41–43). Previous studies established broad-spectrum antiviral activities of SERMs against a number of human pathogens. Interestingly, antiviral activities of SERMs were found to be mostly independent of estrogen receptors and involved other cellular or viral targets (14, 20, 23–25, 44, 45). For example, tamoxifen inhibited the binding and penetration of HSV-1 by targeting the chloride channels (46). A recent study revealed that tamoxifen and clomiphene could suppress the spike protein-induced membrane fusion which is an important early step for severe acute respiratory syndrome coronavirus 2 (SARS-CoV-2) infection (47). Endolysosomal calcium accumulation upon treatment with SERMs obstructed cellular entry by the Ebola virus (45). On the other hand, tamoxifen diminished VSV infection wherein the inhibition was mediated by type-1 interferon (25). Interestingly in flaviviruses, the target for SERMs are the RNA translation and/or replication of the virus (20). In this study anti-CHIKV studies were performed using Vero cells, which lack the expression of estrogen receptor α and produce only a low level of estrogen receptor β (23). Of note, Vero cells also do not secrete type-1 interferon. However, this study has not ruled out that SERMs might influence other host pathways or target other viral proteins of the alphaviruses.

In our mouse infection studies, the therapeutic application of tamoxifen efficiently restricted CHIKV propagation and persistence *in vivo.* We further noticed a modest but significant improvement of the footpad swelling in CHIKV-infected mice administered with a moderate dose of tamoxifen (20 mg/kg). Local inflammation triggered upon CHIKV infection orchestrates the influx of leukocytes, including macrophages and CD4 T cells, in the infected tissue leading to musculoskeletal pathologies (29). Importantly, alphaviruses engage the cellular NF-κB signaling pathway, which promotes inflammatory gene expressions (48, 49). It has been recently shown that besides promoting inflammation, NF-κB signaling also augments replication of SINV in infected neurons (49). Curiously, tamoxifen, independent of estrogen receptor signaling, also potently suppresses the NF-κB pathway and restrains inflammation (36, 50). It is also possible that SERMs relied, at least in part, on the down-modulation of NF-κB signaling for mitigating alphavirus virus-induced disease pathologies. Taken together, we argue that the therapeutic effect of tamoxifen observed in our preclinical model could have been attributed to its both anti-nsP1 activity and generic anti-inflammatory functions. Though we have demonstrated nsP1 inhibition by SERM compounds as a probable mechanism for alphaviral inhibition by the compounds, further studies are warranted to fully dissect the contribution of host factors and viral proteins in generating anti-alphaviral effects of SERMs.

While our results assure that adaptive mutations may necessarily not pose a threat to the translation potential of SERM regimes, future studies ought to more extensively characterize the SERM-alphavirus interaction interface. There is a possibility that adaptive mutations at multiple sites are required to circumvent the inhibitory effect of the compounds. Though our *in vitro* enzymatic assays confirmed inhibition of enzymatic activity of purified CHIKV and VEEV nsP1 proteins, computational analysis of protein-ligand binding (including extensive molecular dynamic simulations) and site-directed mutagenesis combined with *in vitro* activity studies will be required to accurately assess the nsP1-SERM interaction and elucidate the binding preferences of similar compounds to the alphaviral capping enzyme. Nonetheless, our current work offers a proof of concept that SERM compounds can act as direct acting antivirals by impairment of alphaviral nsP1 thereby providing another estrogen receptor signalling-independent mechanism for antiviral effect of SERMs.

In conclusion, this report identifies and characterizes SERMs as promising lead candidates for further development of targeted anti-alphaviral therapy. Dose-dependent inhibition of CHIKV and SINV by all three tested SERMs, and inhibition of CHIKV nsP1 and VEEV nsP1 enzymatic activity affirms the broad-spectrum antiviral activity of SERMs against alphaviruses. A number of SERMs are FDA-approved, have good oral availability, established pharmacological and safety profiles, and are already in clinical practice for half a century. With the effective inhibition of alphaviral replication in cell culture and mice model by SERM compounds, a possible new approach of using SERMs and their superior analogs can be further investigated for the identification and development of pan-alphavirus therapeutic agents.

## Acknowledgements

We thank Dr. Richard Kuhn at Purdue University, USA for providing SINV-FLuc plasmid. R.M., S.C., and M.A. are thankful to council of scientific and industrial research, India. A.D. thanks Department of Biotechnology, India (DBT), and Y.R. thanks Indian Council for Medical Research (ICMR), Govt. of India for research fellowship. Authors thank Biotech Consortium India Limited, DBT, Ministry of Science and Technology, Government of India, New Delhi, India for their assistance in filing the patent for “An antiviral composition against alphaviruses” (Patent file no. 201911049130, dated 29th November 2019).

## Transparency declarations

None to declare.

## References

1. Mostafavi H, Abeyratne E, Zaid A, Taylor A. 2019. Arthritogenic Alphavirus-Induced Immunopathology and Targeting Host Inflammation as A Therapeutic Strategy for Alphaviral Disease. 3. Viruses 11:290.

2. Strauss JH, Strauss EG. 1994. The alphaviruses: gene expression, replication, and evolution. Microbiol Rev 58:491–562.

3. Pialoux G, Gaüzère B-A, Jauréguiberry S, Strobel M. 2007. Chikungunya, an epidemic arbovirosis. Lancet Infect Dis 7:319–327.

4. Soumahoro M-K, Boelle P-Y, Gaüzere B-A, Atsou K, Pelat C, Lambert B, Ruche GL, Gastellu-Etchegorry M, Renault P, Sarazin M, Yazdanpanah Y, Flahault A, Malvy D, Hanslik T. 2011. The Chikungunya Epidemic on La Réunion Island in 2005–2006: A Cost-of-Illness Study. PLoS Negl Trop Dis 5:e1197.

5. Adouchief S, Smura T, Sane J, Vapalahti O, Kurkela S. 2016. Sindbis virus as a human pathogen-epidemiology, clinical picture and pathogenesis. Rev Med Virol 26:221–241.

6. Strauss EG, Rice CM, Strauss JH. 1984. Complete nucleotide sequence of the genomic RNA of Sindbis virus. Virology 133:92–110.

7. Rupp JC, Sokoloski KJ, Gebhart NN, Hardy RW. 2015. Alphavirus RNA synthesis and non-structural protein functions. J Gen Virol 96:2483–2500.

8. Ahola T, Kääriäinen L. 1995. Reaction in alphavirus mRNA capping: formation of a covalent complex of nonstructural protein nsP1 with 7-methyl-GMP. Proc Natl Acad Sci 92:507–511.

9. Hyde JL, Diamond MS. 2015. Innate immune restriction and antagonism of viral RNA lacking 2’-O methylation. Virology 0:66–74.

10. Tomar S, Narwal M, Harms E, Smith JL, Kuhn RJ. 2011. Heterologous production, purification and characterization of enzymatically active Sindbis virus nonstructural protein nsP1. Protein Expr Purif 79:277–284.

11. Feibelman KM, Fuller BP, Li L, LaBarbera DV, Geiss BJ. 2018. Identification of small molecule inhibitors of the Chikungunya virus nsP1 RNA capping enzyme. Antiviral Res 154:124–131.

12. Inhibition of Chikungunya virus by an Adenosine Analog targeting the SAM dependent nsP1 methyltransferase – Mudgal -- FEBS Letters – Wiley Online Library.

13. Pushpakom S, Iorio F, Eyers PA, Escott KJ, Hopper S, Wells A, Doig A, Guilliams T, Latimer J, McNamee C, Norris A, Sanseau P, Cavalla D, Pirmohamed M. 2019. Drug repurposing: progress, challenges and recommendations. 1. Nat Rev Drug Discov 18:41–58.

14. Montoya MC, Krysan DJ. 2018. Repurposing Estrogen Receptor Antagonists for the Treatment of Infectious Disease. mBio 9:e02272–18.

15. Plouffe L. 2000. Selective estrogen receptor modulators (SERMs) in clinical practice. J Soc Gynecol Investig 7:S38–46.

16. Legha SS. 1988. Tamoxifen in the Treatment of Breast Cancer. Ann Intern Med 109:219–228.

17. Squirewell EJ, Qin X, Duffel MW. 2014. Endoxifen and Other Metabolites of Tamoxifen Inhibit Human Hydroxysteroid Sulfotransferase 2A1 (hSULT2A1). Drug Metab Dispos 42:1843–1850.

18. Homburg R. 2005. Clomiphene citrate—end of an era? a mini-review. Hum Reprod 20:2043–2051.

19. Wibowo E, Pollock PA, Hollis N, Wassersug RJ. 2016. Tamoxifen in men: a review of adverse events. Andrology 4:776–788.

20. Eyre NS, Kirby EN, Anfiteatro DR, Bracho G, Russo AG, White PA, Aloia AL, Beard MR. 2020. Identification of Estrogen Receptor Modulators as Inhibitors of Flavivirus Infection. Antimicrob Agents Chemother 64:e00289-20, /aac/64/8/AAC.00289-20.atom.

21. Watashi K, Inoue D, Hijikata M, Goto K, Aly HH, Shimotohno K. 2007. Anti-hepatitis C virus activity of tamoxifen reveals the functional association of estrogen receptor with viral RNA polymerase NS5B. J Biol Chem 282:32765–32772.

22. Tohma D, Tajima S, Kato F, Sato H, Kakisaka M, Hishiki T, Kataoka M, Takeyama H, Lim C-K, Aida Y, Saijo M. 2019. An estrogen antagonist, cyclofenil, has anti-dengue-virus activity. Arch Virol 164:225–234.

23. Johansen LM, Brannan JM, Delos SE, Shoemaker CJ, Stossel A, Lear C, Hoffstrom BG, DeWald LE, Schornberg KL, Scully C. 2013. FDA-approved selective estrogen receptor modulators inhibit Ebola virus infection. Sci Transl Med 5:190ra79–190ra79.

24. Zheng K, Chen M, Xiang Y, Ma K, Jin F, Wang X, Wang X, Wang S, Wang Y. 2014. Inhibition of herpes simplex virus type 1 entry by chloride channel inhibitors tamoxifen and NPPB. Biochem Biophys Res Commun 446:990–996.

25. Cham LB, Friedrich S-K, Adomati T, Bhat H, Schiller M, Bergerhausen M, Hamdan T, Li F, Machlah YM, Ali M, Duhan V, Lang KS, Friebus-Kardash J, Lang J. 2019. Tamoxifen Protects from Vesicular Stomatitis Virus Infection. Pharmaceuticals 12:142.

26. Singh H, Mudgal R, Narwal M, Kaur R, Singh VA, Malik A, Chaudhary M, Tomar S. 2018. Chikungunya virus inhibition by peptidomimetic inhibitors targeting virus-specific cysteine protease. Biochimie 149:51–61.

27. Kaur R, Neetu null, Mudgal R, Jose J, Kumar P, Tomar S. 2019. Glycan-dependent chikungunya viral infection divulged by antiviral activity of NAG specific chi-like lectin. Virology 526:91–98.

28. Kaur R, Mudgal R, Narwal M, Tomar S. 2018. Development of an ELISA assay for screening inhibitors against divalent metal ion dependent alphavirus capping enzyme. Virus Res https://doi.org/10.1016/j.virusres.2018.06.013.

29. Gardner J, Anraku I, Le TT, Larcher T, Major L, Roques P, Schroder WA, Higgs S, Suhrbier A. 2010. Chikungunya virus arthritis in adult wild-type mice. J Virol 84:8021–8032.

30. Teo T-H, Chan Y-H, Lee WWL, Lum F-M, Amrun SN, Her Z, Rajarethinam R, Merits A, Rötzschke O, Rénia L, Ng LFP. 2017. Fingolimod treatment abrogates chikungunya virus– induced arthralgia. Sci Transl Med 9.

31. Agarwal A, Singh AK, Sharma S, Soni M, Thakur AK, Gopalan N, Parida MM, Rao PVL, Dash PK. 2013. Application of real-time RT-PCR in vector surveillance and assessment of replication kinetics of an emerging novel ECSA genotype of Chikungunya virus in Aedes aegypti. J Virol Methods 193:419–425.

32. Sourisseau M, Schilte C, Casartelli N, Trouillet C, Guivel-Benhassine F, Rudnicka D, Sol-Foulon N, Roux KL, Prevost M-C, Fsihi H, Frenkiel M-P, Blanchet F, Afonso PV, Ceccaldi P-E, Ozden S, Gessain A, Schuffenecker I, Verhasselt B, Zamborlini A, Saïb A, Rey FA, Arenzana-Seisdedos F, Desprès P, Michault A, Albert ML, Schwartz O. 2007. Characterization of Reemerging Chikungunya Virus. PLoS Pathog 3.

33. Rana J, Rajasekharan S, Gulati S, Dudha N, Gupta A, Chaudhary VK, Gupta S. 2014. Network mapping among the functional domains of Chikungunya virus nonstructural proteins. Proteins 82:2403–2411.

34. Sreejith R, Rana J, Dudha N, Kumar K, Gabrani R, Sharma SK, Gupta A, Vrati S, Chaudhary VK, Gupta S. 2012. Mapping interactions of Chikungunya virus nonstructural proteins. Virus Res 169:231–236.

35. Žusinaite E, Tints K, Kiiver K, Spuul P, Karo-Astover L, Merits A, Sarand I. 2007. Mutations at the palmitoylation site of non-structural protein nsP1 of Semliki Forest virus attenuate virus replication and cause accumulation of compensatory mutations. J Gen Virol 88:1977–1985.

36. Takada Y, Bhardwaj A, Potdar P, Aggarwal BB. 2004. Nonsteroidal anti-inflammatory agents differ in their ability to suppress NF-κB activation, inhibition of expression of cyclooxygenase-2 and cyclin D1, and abrogation of tumor cell proliferation. Oncogene 23:9247–9258.

37. Rice CM, Levis R, Strauss JH, Huang HV. 1987. Production of infectious RNA transcripts from Sindbis virus cDNA clones: mapping of lethal mutations, rescue of a temperature-sensitive marker, and in vitro mutagenesis to generate defined mutants. J Virol 61:3809–3819.

38. Sokoloski KJ, Haist KC, Morrison TE, Mukhopadhyay S, Hardy RW. 2015. Noncapped Alphavirus Genomic RNAs and Their Role during Infection. J Virol 89:6080–6092.

39. Wang Y-F, Sawicki SG, Sawicki DL. 1994. Alphavirus nsP3 functions to form replication complexes transcribing negative-strand RNA. 10. J Virol 68:6466–6475.

40. Wang YF, Sawicki SG, Sawicki DL. 1991. Sindbis virus nsP1 functions in negative-strand RNA synthesis. J Virol 65:985–988.

41. Wiseman H, Laughton MJ, Arnstein HR, Cannon M, Halliwell B. 1990. The antioxidant action of tamoxifen and its metabolites.Inhibition of lipid peroxidation. FEBS Lett 263:192–194.

42. O’Brian CA, Housey GM, Weinstein IB. Specific and Direct Binding of Protein Kinase C to an Immobilized Tamoxifen Analogue 5.

43. Tamoxifen inhibits acidification in cells independent of the estrogen receptor | PNAS.

44. Murakami Y, Fukasawa M, Kaneko Y, Suzuki T, Wakita T, Fukazawa H. 2013. Selective estrogen receptor modulators inhibit hepatitis C virus infection at multiple steps of the virus life cycle. Microbes Infect 15:45–55.

45. Fan H, Du X, Zhang J, Zheng H, Lu X, Wu Q, Li H, Wang H, Shi Y, Gao G, Zhou Z, Tan D-X, Li X. 2017. Selective inhibition of Ebola entry with selective estrogen receptor modulators by disrupting the endolysosomal calcium. 1. Sci Rep 7:41226.

46. Zheng K, Chen M, Xiang Y, Ma K, Jin F, Wang X, Wang X, Wang S, Wang Y. 2014. Inhibition of herpes simplex virus type 1 entry by chloride channel inhibitors tamoxifen and NPPB. 4. Biochem Biophys Res Commun 446:990–996.

47. Zu S, Luo D, Li L, Ye Q, Li R-T, Wang Y, Gao M, Yang H, Deng Y-Q, Cheng G. 2021. Tamoxifen and clomiphene inhibit SARS-CoV-2 infection by suppressing viral entry. Signal Transduct Target Ther 6:1–4.

48. Bais SS, Ratra Y, Khan NA, Pandey R, Kushawaha PK, Tomar S, Medigeshi G, Singh A, Basak S. Chandipura Virus Utilizes the Prosurvival Function of RelA NF-κB for Its Propagation. J Virol 93:e00081–19.

49. Yeh JX, Park E, Schultz KLW, Griffin DE. NF-κB Activation Promotes Alphavirus Replication in Mature Neurons. J Virol 93:e01071–19.

50. Behjati S, Frank MH. 2009. The effects of tamoxifen on immunity. Curr Med Chem 16:3076–3080.

