## Supplemental Data for "Selective estrogen receptor modulators limit alphavirus infection by targeting the viral capping enzyme nsP1"

**SUPPLEMENTARY INFORMATION**

**MTT and LDH assay:** 96-well plates were seeded with Vero cells at 1x10^4^ cells/well and incubated for 24 h at 37°C and 5% CO_2_. Media was removed from subconfluent monolayer and the compound dilutions prepared in serum-free media were added to each well. After 24 h, 20 μl of 5 mg/ml MTT was added to each well and incubated for 4 h at 37°C. The absorbance was then measured at 570 nm and 630 nm with the Cytation 3 multi-mode plate reader after the solubilisation of reduced formazan in the cells with DMSO (BioTek Instruments, Inc.). Lactate Dehydrogenase assay (LDH) was carried out using an LDH assay kit (Thermo Fisher Scientific, USA), according to manufacturer’s instructions, to evaluate the cytotoxicity of various compounds. The CC_50_ values were calculated using Graph pad’s non-linear regression curve fit.

**Luciferase-based antiviral assay**

The luciferase-based assay used a genetically engineered strain of SINV (SINV-FLuc) wherein the *firefly luciferase* gene is driven by an internal ribosome entry site element (IRES) ^1,2^. A reporter SINV replicon (SINV-REP) was constructed in which all the structural proteins downstream of subgenomic (SG) promoter were replaced with FLuc and IRES element at the 3´end.

BHK-21 cells were electroporated in a 0.2 cm cuvette (25 μF, 1.5 kV, and 200 Ω) with 10 µg of capped RNA transcripts for SINV-FLuc or SINV-REP. For SINV-REP, 4 h post-transfection, compounds or 0.1% DMSO were added to the transfected cells, and cell lysate was collected 6 h after the addition of compounds. For SINV-FLuc, media was collected 48 h post-transfection and assayed for titer determination. For inhibition assays, BHK-21 cells at a confluency of 80-90% in a 24-well plate were infected with SINV-FLuc at MOI of 0.1 for 1.5 h. Media containing compounds or 0.1% DMSO was added to the infected cells and lysate was collected at 12 hpi. Luciferase activity of lysates was measured using a firefly luciferase assay kit (Promega, USA) according to the manufacturer’s instructions.

**Conventional plaque assay:** Vero cells were plated in 24-well plate and incubated till cells reached 90% confluency. Ten-fold serial dilutions of supernatants collected from virus-infected cells were prepared. Vero cells were incubated with the dilutions. After 1.5 h, the media was removed and the cells were covered with the overlay media containing 1:1 ratio of 2X MEM (with 5% FBS) and carboxymethyl cellulose (2%). The cells were then incubated for 48 h and then the overlay media was removed. Cells were fixed using 10% formaldehyde and incubated overnight in the dark at room temperature. For visualization of plaques, the cells were stained with 1% crystal violet. The numbers of plaques were counted and PFU/ml (plaque-forming units per ml) was calculated using the formula -

PFU/ml = (No. of Plaques)/ (Dilution factor x Volume of diluted virus per well)

**qRT-PCR:** qRT-PCR assay was performed to quantify intracellular viral RNA or those present in samples derived from infected mice. CHIKV infected cells were treated with 4-OHT, clomifene, or tamoxifen with MOI:1. To quantify intracellular RNA at 6 h, Vero cells were infected with CHIKV or SINV at an MOI of 5 and infected cells were treated with compounds. Total RNA was isolated from the cells 6 h post-infection (hpi) or 24 hpi using Trizol following manufacturer’s instructions. Subsequently, cDNA was synthesized using Accuscript High-fidelity 1^st^ strand cDNA synthesis kit (Agilent technologies, USA) in accordance with manufacturer’s protocol. Actin RNA and the viral RNA was quantified using KAPA SYBR fast universal qPCR kit (Sigma-Aldrich, USA) on Quantstudio real-time PCR system (Applied Biosystems, CA) following manufacturer’s protocol. The amplified products were verified by melting curve analysis. The experiment was performed in triplicate and ΔΔCt method was used to calculate relative values.

The primer sequences used were as follows:

CHIKV E1 forward: 5′-AAGTACACTGTGCAGCTGAGT-3′;

CHIKV E1 reverse: 5′-GCATAGCACCACGATTAGAATC-3';

SINV E1 forward: 5′-ACAGCCAGATGAGTGAGGCGTA-3′;

SINV E1 reverse: 5′-CGATGCTGAAATTGGTCCAGCTATG-3′;

Actin forward: 5′-ATTGCCGACAGGATGCAGAA-3′, and

Actin reverse: 5′-GCTGATCCACATCTGCTGGAA-3'

**Extracellular vRNA quantitation:**

The quantitative analysis of the extracellular SINV RNA was accomplished as described previously.^3^ Vero cells were infected with SINV at an MOI of 1 and the infected cells were treated with 2.5 μM and 5 μM concentration of compounds. Supernatant was collected 24 hpi and 5 μl of supernatant was reverse transcribed using SINV E1 primers as described above. A standard curve for viral RNA was generated and was used to determine absolute quantities of RNA.

**Selection of resistant mutants:** Subconfluent monolayer of Vero cells was infected with CHIKV at an MOI of 1 and cells were then treated with 2.5 μM of compounds for 5 passages, 5 μM concentration for the next 5 passages, and inhibitor concentration was increased to 7.5 μM for another 5 passages. Viral-induced CPE relative to mock-treated cells were monitored every 12 h with a light inverted microscope (Carl Zeiss, Germany) to confirm the development of resistant variants of the viruses. Media was harvested upon the appearance of CPE in the infected cells at each passage. Viral titers were determined after every passage and viral stocks were diluted to infect the cells in the subsequent passage at an MOI of 1. After passaging the CHIKV-infected cells in the presence of 4-OHT, partially-resistant mutants against 4-OHT were plaque-purified and used to infect Vero cells in the presence 7.5 μM 4-OHT. vRNA was isolated 24 hpi and cDNA generation was performed from harvested supernatant using random hexamers using Accuscript High-fidelity 1^st^ strand cDNA synthesis kit (Agilent technologies, USA). Resistant genotype was determined by next generation sequencing. Nucleotide sequence of the clinical isolate of CHIKV (strain no.: 119067; GenBank: KY057363.1) was used for sequence comparison.^4^

**Immunofluorescence assay (IFA):** The antiviral activity of various compounds was tested using indirect IFA. Briefly, subconfluent monolayer of Vero cells in a 6-well plate was infected with CHIKV at MOI 1 for 1.5 h, then the compound dilutions (in maintenance media) were added and cells were incubated for 36 h. Cells were then fixed using 1:1 ratio of methanol and acetone for 30 min at -20⁰C. 500 μl of 0.5% Triton X-100 was added to each well and incubated for 7 min at room temperature to facilitate permeabilization of cells. Cells were washed three times between two consecutive steps with PBS. Cells were then incubated with 100 μl/well of primary antibody (1:250; Santa Cruz Biotechnology Inc.) against CHIKV for 1 h at 37⁰C. Cells were further washed and incubated with 100 μl fluorescein isothiocyanate (FITC)-conjugated anti-mouse secondary antibody (1:500; Sigma) for 1 h. Finally, cells were counter-stained using 4',6-diamidino-2-phenylindole (DAPI; Himedia, India), incubated for 15 min, and then observed under EVOS FL imaging system (Thermo Fisher Scientific, USA) using both DAPI and GFP channels. EVOS FL software was used to process acquired images.

**Time-of-addition assay:** Vero cell monolayer in a 24-well plate was infected with CHIKV or SINV viruses at MOI of 1. Cells were treated with 0.1% DMSO, or 5 µM of 4-OHT, or 7.5 µM tamoxifen/clomifene. For pretreatment assay, cells were incubated with compounds 2 h prior to infection with the virus. Media containing compounds was removed and cells were washed twice with PBS before adding viral inoculums and subsequently low-serum maintenance media was added post-infection. For the cotreatment assay, compounds were added to the monolayer simultaneously with the virus inoculum and monolayer was washed post-infection before the addition of maintenance media. Alternatively, for posttreatment assay, compounds were added at four different time points (0, 2, 4, and 6 h) to the infected cells. At 24 hpi, culture supernatants were harvested for viral titer determination using a conventional plaque-forming assay.

**TCID_50_ methods:** Briefly, serum or skeletal muscle extracts were serially diluted in DMEM supplemented with 2% FBS. Four wells in a 96 well plate were designated for each dilution and 100 μl of the each dilution was transferred per well. Each well was then seeded with 10^4^ Vero cells and cells were cultured for another three to four days. Uninfected cells were used as control. Next, media was discarded and 100 μl of 10% formaldehyde (Sigma) were added per well to fix the cells. The plate was incubated for six hours in the dark at room temperature. After fixation, formaldehyde solution was discarded and 50 μl of crystal violet solution (0.2% crystal violet dissolved in 2% ethanol) was added per well and incubated for another 5 minutes. Then, a single wash for 30 seconds after incubation was given and cytopathic effects were analyzed. As described, the dilution at which 50% of cells were infected was determined and reported as log_10_ (TCID_50_/gm).^5^

**REFERENCES**

1. Snyder JE, Kulcsar KA, Schultz KL, *et al.* Functional characterization of the alphavirus TF protein. *J Virol* 2013: JVI-00449.

2. Kaur R, Mudgal R, Narwal M, Tomar S. Development of an ELISA assay for screening inhibitors against divalent metal ion dependent alphavirus capping enzyme. *Virus Res* 2018. Available at: http://www.sciencedirect.com/science/article/pii/S0168170218300285. Accessed September 1, 2018.

3. Sokoloski KJ, Hayes CA, Dunn MP, Balke JL, Hardy RW, Mukhopadhyay S. Sindbis virus infectivity improves during the course of infection in both mammalian and mosquito cells. *Virus Res* 2012; **167**: 26–33.

4. Singh H, Mudgal R, Narwal M, *et al.* Chikungunya virus inhibition by peptidomimetic inhibitors targeting virus-specific cysteine protease. *Biochimie* 2018; **149**: 51–61.

5. Sourisseau M, Schilte C, Casartelli N, *et al.* Characterization of Reemerging Chikungunya Virus. *PLoS Pathog* 2007; **3**. Available at: https://www.ncbi.nlm.nih.gov/pmc/articles/PMC1904475/. Accessed April 7, 2021.

**Supplementary Figures:**

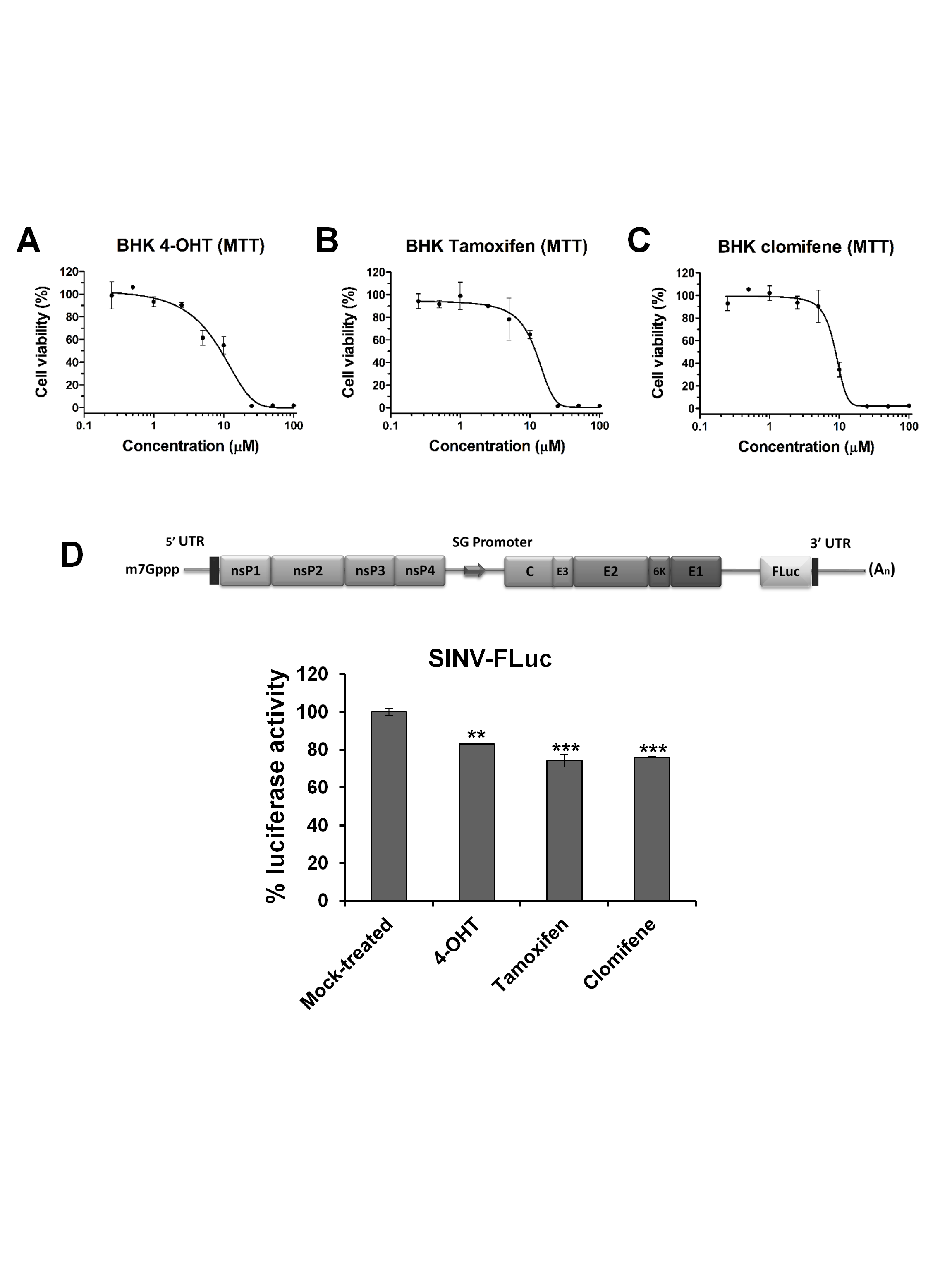

**Figure S1: Viability of BHK-21 cells.** Viability of BHK-21 cells as determined by standard MTT assay after 24 h of incubation at indicated doses of (A) 4-OHT, (B) tamoxifen, and (C) clomifene. 0.1% DMSO was used as a positive control. The experiments were done in triplicates. Data were normalized using Graph pad’s non-linear regression curve fit and the calculated 50% cytotoxic concentration (CC_50_) values are summarized in Table 1. Values are mean and error bars are Standard Deviation.

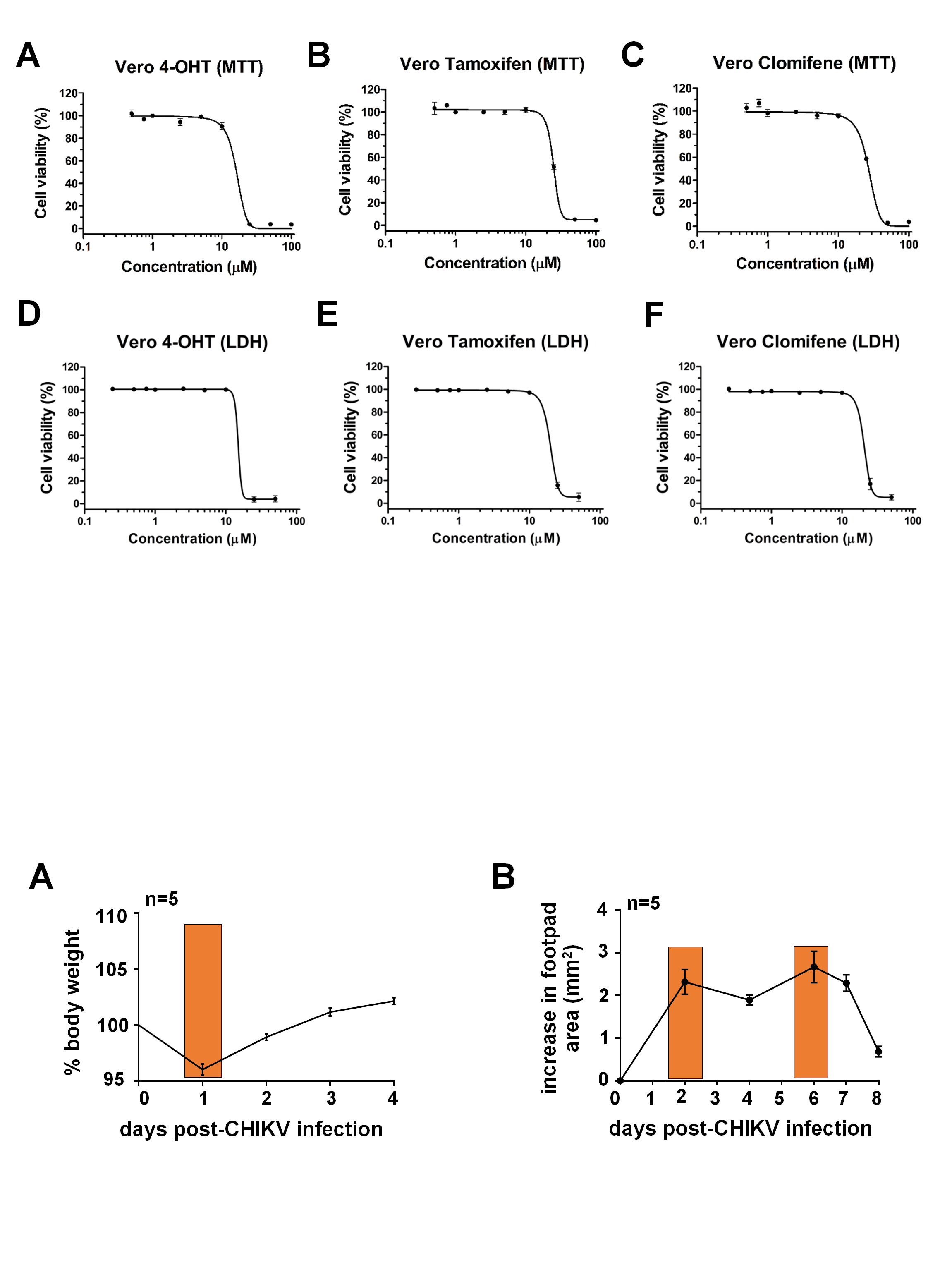

**Figure S2:** **Viability of Vero cells as determined by MTT and LDH assay.** (A-C) MTT cytotoxicity assay for Vero cells treated with different concentrations of SERMs. Graphs represent percent viability of Vero cells on treatment with (A) 4-OHT, (B) tamoxifen, and (C) clomifene. Survival dose-response assay with LDH kit for (D) 4-OHT, (E) tamoxifen, and (F) clomifene. 0.1% DMSO was used as a positive control. Values are mean and error bars are standard deviation of three biological replicates. Data were normalized using Graph pad’s non-linear regression curve fit and the calculated 50% cytotoxic concentration (CC_50_) values are summarized in Table 1.

**Figure S3: Dose-response inhibition assay for 4-OHT-resistant CHIKV variant.** Vero cells were infected with either wild-type CHIKV or 4-OHT-resistant variant at an MOI of 1 in the presence of indicated concentrations of 4-OHT. Media was harvested 24 hpi. Viral titers were determined by plaque assay on Vero cells. Viral titer in corresponding vehicle-treated cells was considered as 100%. Data were normalized using Graph pad’s non-linear regression curve fit and 50% effective concentration (EC_50_) values were calculated. Values are mean from three independent experiments and error bars are standard deviation.

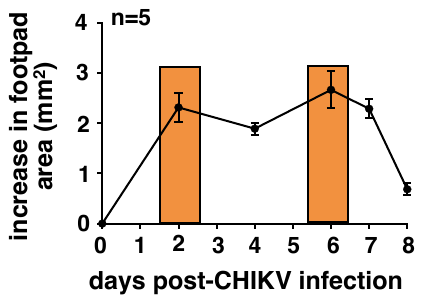

**Figure S4: Examining CHIKV infected mice:** Time course analyses revealing changes in the footpad area of WT C57BL/6 mice upon infection with 10^6^ PFU of CHIKV. Shaded areas represent the time points further examined for tamoxifen treatment.
